# IL-12 Secreting CAR-T Cells Reprogram the Tumor Microenvironment and Improve Efficacy Against Heterogeneous Models of Glioblastoma

**DOI:** 10.1101/2025.06.04.657937

**Authors:** Steven H. Shen, Aditya A. Mohan, Kelly M. Hotchkiss, Sarah L. Cook, Kisha K. Patel, Eliese Moelker, Adam M. Swartz, Carter M. Suryadevara, Daniel Wilkinson, Peter E Fecci, Luis Sanchez-Perez, John H. Sampson, Anoop P. Patel

## Abstract

**Background:** Glioblastoma (GBM) remains uniformly lethal due to pronounced intratumoral heterogeneity and a highly immunosuppressive microenvironment that limits the efficacy of targeted therapies.

**Methods:** We engineered chimeric antigen receptor (CAR) T cells targeting EGFRvIII and armored them with a single-chain interleukin-12 (scIL12) payload. These cells were tested in syngeneic, orthotopic GBM mouse models exhibiting heterogeneous EGFRvIII expression. CAR T cells were delivered intracranially without lymphodepletion.

**Results:** Intracranial administration of of scIL12-secreting CAR-T cells eradicated tumors without requiring lymphodepletion, achieving 50% long-term survival. Survival benefits depended entirely on endogenous CD8⁺ T cells, as efficacy was abolished in CD8-deficient hosts and unaffected by NK cell depletion. Notably, therapeutic efficacy was abrogated by lymphodepletion, underscoring the necessity of an intact endogenous immune response. Mechanistically, scIL12 enhanced CAR-T cell persistence and reprogrammed tumor-associated microglia, indicating potential epitope spreading through polyclonal endogenous CD8+ T-cell responses, which facilitate the elimination of EGFRvIII-negative tumor cells.

**Conclusions:** This study demonstrates the pleiotropic benefits of IL-12 armored CAR-T cells with improved targeting of antigen-positive tumor cells and simultaneous remodeling of the microenvironment to engage adaptive immunity against antigen-negative clones. This strategy offers a potential clinically actionable approach to improve outcomes in GBM by circumventing the need for toxic lymphodepletion and addressing tumor heterogeneity.

## INTRODUCTION

Glioblastoma (GBM) is the most common and lethal primary malignant brain tumor in adults. Despite aggressive multimodal therapy, including maximal safe resection, radiotherapy, and temozolomide, median survival remains just 15–20 months (1, 2). Clinical trials using both targeted therapy and immunotherapy, including immune checkpoint blockade and adoptive T cell therapies, have consistently failed to improve outcomes (3) (4). A central driver of this therapeutic resistance is intratumoral heterogeneity—a hallmark of GBM that enables tumor subpopulations to evade single-target therapies through antigen loss or transcriptional plasticity (5) (6). Single-cell RNA sequencing (scRNA-seq) studies have systematically mapped this heterogeneity, revealing dynamic shifts in oncogenic signaling, metabolic states, and immune evasion mechanisms across tumor cells and the microenvironment (7). Consequently, therapies targeting highly expressed but heterogeneous antigens have not demonstrated clinical benefits (8), highlighting the need for therapeutic strategies specifically designed to address tumor heterogeneity.

EGFRvIII, a tumor-specific neoantigen generated by an in-frame deletion of exons 2–7 in *EGFR*, has long been a focus of immunotherapy development due to exquisite tumor specificity (9). Early clinical trials of EGFRvIII-targeted vaccines and chimeric antigen receptor (CAR) T cells showed transient responses, but long-term efficacy has historically been limited by heterogeneous antigen expression (8). This underscores a critical need for immunotherapeutic strategies that target both antigen-positive cells and engage endogenous immunity to eliminate antigen-negative populations through epitope spreading, a process wherein tumor antigens released by dying cells are cross-presented to T cells to enable a broad anti-tumor T cell response (10).

A major barrier to epitope spreading in GBM is the reliance of conventional CAR-T therapies on lymphodepletion to enhance engraftment by depleting regulatory T cells (Tregs) and increasing availability of homeostatic cytokines like interleukin-7 (IL-7) and interleukin-15 (IL-15)(11). While lymphodepletion transiently enhances CAR-T cell persistence, it also depletes endogenous immune effectors required for epitope spreading (10). To address this paradox, we hypothesized that “armored” CAR-T cells engineered to secrete immunostimulatory/homeostatic cytokines could obviate the need for lymphodepletion while augmenting CAR-T persistence and efficacy. Interleukin-12 (IL-12), a pleiotropic cytokine, enhances T cell cytotoxicity, promotes dendritic cell (DC) maturation, and reprograms immunosuppressive myeloid populations (12–14). In preclinical studies, the combination of intratumoral delivery of IL-12 fused to the Fc domain of IgG3, low-dose total body irradiation, and intravenous CAR-T cell administration has shown success in treating EGFRvIII-expressing GL261 intracranial tumors. IL-12 was found to reinvigorate dysfunctional CAR-T cells, prevent T cell exhaustion, and remodel the myeloid compartment(15). However, the question of whether IL-12 can synergize with CAR-T cell therapy to overcome antigen heterogeneity in immunocompetent models remains unexplored.

To that end, we report the development of EGFRvIII-targeted CAR-T cells armored with single-chain IL-12 (scIL12) that extends survival in a syngeneic, antigenically heterogeneous model of GBM without the need for lymphodepletion. Using an immunocompetent, orthotopic mouse model of GBM that contains a mixture of cells that are antigen positive and negative, we show that scIL12 secretion enables CAR-T cells to persist in the TME, reprogram immunosuppressive microglia, and recruit endogenous CD8+ T cells to eliminate antigen-negative tumor cells. Our findings establish EGFRvIII CAR scIL12 T cells as a multifaceted therapeutic strategy capable of addressing the dual challenges of antigen heterogeneity and immunosuppression while obviating the need for toxic lymphodepletion regimens.

## RESULTS

### Generation and In Vitro Validation of IL-12 Armored CAR-T cells

We previously developed a retroviral vector encoding a third-generation chimeric antigen receptor (CAR) targeting the EGFRvIII antigen based on the mAb139 monoclonal antibody (mAb139)(16) (Fig. 1A-B). Given the extensive literature supporting IL-12 as an armored CAR-T cell strategy for enhancing anti-tumor immunity, we engineered an enhanced CAR vector incorporating an encephalomyocarditis virus (EMCV) internal ribosomal entry site (IRES) followed by a fusion construct encoding IL-12 p35 and p40 subunits, connected by a flexible linker (Fig. 1A-B). Flow cytometry confirmed efficient retroviral transduction of both standard EGFRvIII CAR and IL-12 armored EGFRvIII CAR (EGFRvIII CAR scIL12) constructs (Fig. 1C), and although there was a slight reduction in the proportion of cells that were transduced with EGFRvIII CAR scIL12, probably due to the 2233 bp increase in vector size, this difference was not significant (Fig. 1D-E). ELISA assay experiments further confirmed IL-12 secretion exclusively from EGFRvIII CAR scIL12 transduced cells (Fig. 1F). To assess whether IL-12 secretion affects tumor recognition and cytotoxicity, EGFRvIII CAR, EGFRvIII CAR scIL12, and untransduced control T cells were co-cultured with CFSE-labeled CT2AvIII glioma cells. Within 24 hours, both CAR-T cell groups exhibited robust tumor killing across multiple effector-to-target ratios (Fig.1G) as compared to untransduced controls. Collectively, these data demonstrate the successful generation of EGFRvIII CAR scIL12 T cells that maintain tumor-directed cytotoxicity while producing IL-12.

**Figure 1.**
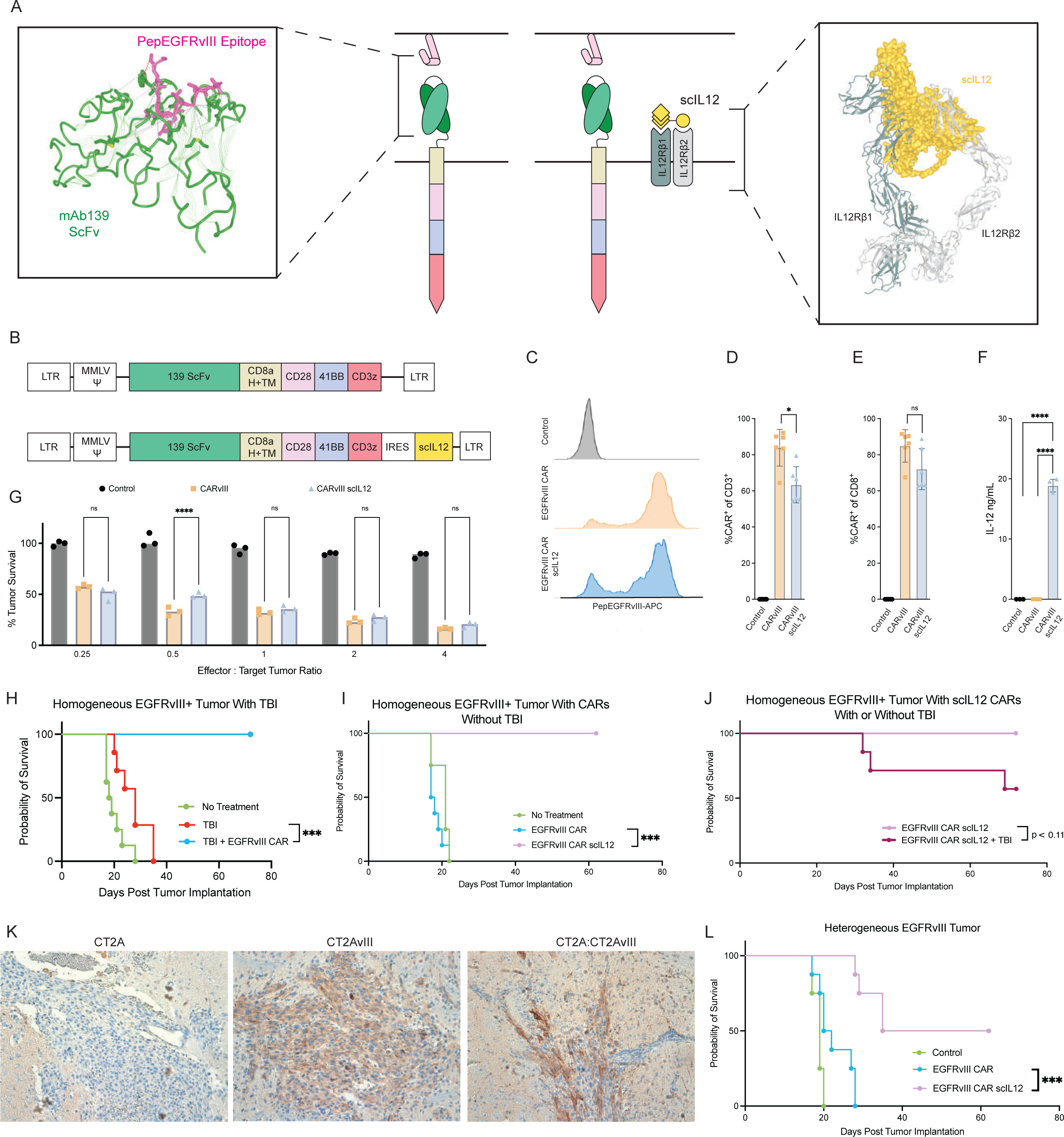
Engineering and Functional Characterization of EGFRvIII CAR scIL12. (A-B) Schematic of the EGFRvIII CAR constructs with and without IL-12 armoring. The third-generation EGFRvIII-specific CAR consists of a mAb139-derived single-chain variable fragment (scFv) for antigen recognition, a CD8α hinge and transmembrane domain, and CD28, 4-1BB, and CD3ζ signaling domains for T cell activation. The IL-12-armored CAR construct includes an internal ribosome entry site (IRES) to drive co-expression of a single-chain IL-12 (scIL12) fusion protein, comprising the IL-12 p35 and IL-12 p40 subunits connected by a flexible linker. The secreted IL-12 binds to IL-12 receptors (IL-12Rβ1/β2) expressed on T cells and myeloid cells to enhance CAR-T cell function and remodel the tumor microenvironment. (C) Flow cytometry histograms confirming EGFRvIII CAR expression in murine CD8^⁺^ T cells. Untransduced T cells (Control), EGFRvIII CAR-T cells, and EGFRvIII CAR scIL12 T cells were stained with APC-labeled PepEGFRvIII peptide multimers. Comparable CAR expression was observed in the EGFRvIII CAR and EGFRvIII CAR scIL12 groups, indicating that IL-12 co-expression does not affect transduction efficiency. (D-E) Quantification of CAR expression in transduced T cells. The proportion of CAR-expressing cells within the CD8^⁺^ and CD3^⁺^ T cell populations was similar between the EGFRvIII CAR and EGFRvIII CAR scIL12 groups, confirming that the addition of IL-12 does not compromise transduction efficiency or CAR stability. (F) IL-12 secretion in CAR-T cell cultures. ELISA of culture supernatants demonstrated that EGFRvIII CAR scIL12 T cells secrete high levels of IL-12, whereas standard CAR-T cells do not (p < 0.0001). This confirms successful expression and secretion of IL-12. (G) CT2A-EGFRvIII glioblastoma cells were co-cultured with either untransduced T cells (Control), EGFRvIII CAR-T cells, or EGFRvIII CAR scIL12 T cells at multiple effector-to-target (E:T) ratios after 24 hours of co-culture. Tumor survival was determined by flow cytometric analysis of the percentage of viable cells remaining within the CFSE+ population. (H) Survival analysis of mice bearing homogeneous EGFRvIII glioblastomas treated with CAR-T cells and lymphodepletion. C57BL/6J mice were implanted with CT2A-EGFRvIII tumors and treated with either total body irradiation (TBI) alone or TBI + EGFRvIII CAR-T cells. Only TBI + EGFRvIII CAR-T cells resulted in tumor clearance and long-term survival (p < 0.001). (I) Survival analysis in mice with homogeneous EGFRvIII tumors treated with CAR-T cells without TBI, with or without IL-12 armoring. Kaplan-Meier survival curves show that EGFRvIII CAR scIL12 T cells significantly enhance survival compared to standard EGFRvIII CAR-T cells (p < 0.001). (J) Survival analysis of CAR-T cells in homogeneous tumors with or without TBI. While EGFRvIII CAR scIL12 T cells led to 100% survival in mice without TBI, only 50% survival was observed when mice were pre-conditioned with TBI (p < 0.11). (K) Immunohistochemistry (IHC) of tumor sections confirming EGFRvIII antigen heterogeneity. Representative IHC images of CT2A parental (EGFRvIII^⁻^), CT2AvIII (EGFRvIII^⁺^), and CT2A:CT2AvIII (mixed) tumors stained for EGFRvIII (brown) and nuclei using hematoxylin (blue). (L) EGFRvIII CAR scIL12 promote durable survival in heterogeneous tumors without TBI. Kaplan-Meier survival analysis shows that EGFRvIII CAR scIL12 T cells significantly improve survival over conventional CAR-T cells (p < 0.001) in antigen-heterogeneous tumors, supporting the role of IL-12 in engaging endogenous immune responses to overcome tumor antigen escape.

### Lymphodepletion is Required for CAR-T cell Efficacy Against Homogeneous Tumor Models

We next evaluated the therapeutic efficacy of EGFRvIII CAR and EGFRvIII CAR-scIL12 T-cells *in vivo* using syngeneic orthotopic glioblastoma models. CT2AvIII cells were implanted into the right basal ganglia of C57BL/6J mice. One week after tumor engraftment, mice were either left untreated or preconditioned with a single 5 Gy dose of total body irradiation (TBI) for lymphodepletion before intratumoral administration of CAR+ T - cells. Consistent with prior findings(17), TBI alone had limited anti-tumor activity, but when combined with EGFRvIII CAR-T cells, there was complete tumor eradication and long-term survival in all treated animals (Fig.1H).

### IL-12 Armored CAR-T cells Eradicate Homogeneous Tumors Without Lymphodepletion

There is evidence that IL-12 has direct homeostatic effects on CAR-T cells(18), potentially obviating the need for lymphodepletion. Given other work suggesting that IL-12 enhances tumor clearance via epitope spreading and recruitment of endogenous immune effectors, we hypothesized that lymphodepletion would be unnecessary and in fact detrimental to the effect of IL-12. To test this, we examined the efficacy of EGFRvIII CAR scIL12 T cells in the CT2A-EGFRvIII model with and without TBI. While EGFRvIII CAR-T cells failed to extend survival without TBI, EGFRvIII CAR-scIL12 T-cells completely eradicated tumors in all treated mice that did not receive TBI (Fig. 1I). Notably, the complete long term survival seen in EGFRvIII CAR scIL12 treated mice was abolished when mice were lymphodepleted prior to treatment (Fig. 1J). These findings highlight the critical role of endogenous immune cells in mediating the efficacy of EGFRvIII CAR scIL12 T-cells.

### IL-12 Armored CAR-T cells Overcome Tumor Antigen Heterogeneity

To model tumor antigen heterogeneity, we implanted a 50:50 mixture of CT2A (EGFRvIII⁻) and CT2A-EGFRvIII (EGFRvIII⁺) cells. Immunohistochemical staining for EGFRvIII confirmed spatially distinct regions of antigen-positive and antigen-negative cells within tumors (Fig. 1K). We then assessed whether IL-12 secretion could improve CAR-T cell efficacy in the heterogeneous tumor setting. 7 days post implantation of our heterogeneous model, mice were treated with EGFRvIII CAR or EGFRvIII CAR scIL12 and monitored for survival. As expected, EGFRvIII CAR-T cells alone were ineffective, failing to extend survival (Fig. 1L). In contrast, IL-12 armored CAR-T cells significantly improved outcomes, with 50% of treated mice achieving long-term survival.

### CAR-T cells Secreting IL-12 Treat Heterogeneous Tumors through Recruitment of Endogenous CD8⁺ T Cells

We next sought to understand the mechanism of antigen negative tumor cell killing by performing selective depletions to isolate the responsible effector cells. Given the known role of IL-12 in enhancing NK cell-mediated anti-tumor responses (19), we first depleted NK cells using pretreatment and maintenance dosing of an anti-NK1.1 antibody (Fig. 2A), which did not affect survival (Fig. 2B). However, the survival benefit of IL-12 armoring was abrogated in RAG1⁻/⁻ mice, suggesting that the efficacy of IL-12 armored CAR-T cells relies on the presence of endogenous T or B cells (Figure 2B). These findings were validated in TCR α ⁻/⁻ mice, further narrowing the effector to the T cell compartment (Supplemental Fig. 1). We then selectively depleted CD4⁺ T cells (Fig. 2C), which did not affect survival (Fig. 2D). However, experiments in CD8⁻/⁻ mice (Fig. 2E) showed complete abrogation of the armoring effect, demonstrating that the survival benefit of IL-12 secreting CAR-T cells was dependent on endogenous CD8^+^ T cells (Fig. 2F).

**Figure 2.**
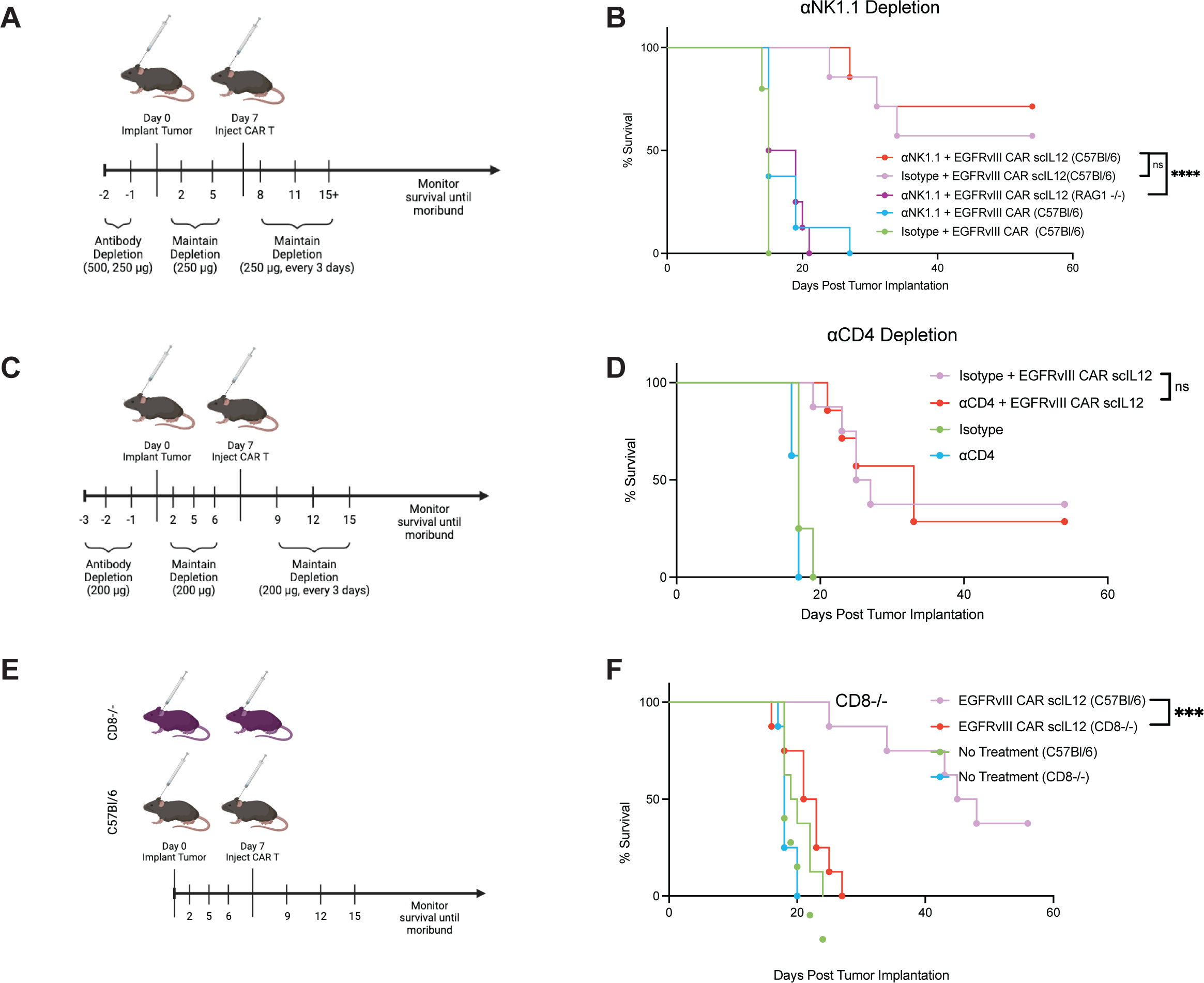
EGFRvIII CAR scIL12Require Endogenous CD8**^⁺^** T Cells but Not NK or CD4**^⁺^** T Cells for Efficacy. (A) Experimental timeline for NK cell depletion studies. Mice were injected with anti-NK1.1 antibodies starting two days before tumor implantation, with maintenance dosing every three days to ensure sustained NK cell depletion. (B) Kaplan-Meier survival analysis evaluating the role of NK cells in IL-12-armored CAR-T cell efficacy. Wild-type C57BL/6J mice and RAG1^⁻^/^⁻^ mice, which lack mature T and B cells but retain functional NK cells, were treated with EGFRvIII CAR scIL12 T cells with or without NK1.1 depletion. Survival outcomes were comparable between NK1.1-depleted and isotype control-treated groups, indicating that NK cells do not play a major role in IL-12-enhanced CAR-T cell therapy (ns, not significant). (C) Experimental timeline for CD4^⁺^ T cell depletion studies. Mice received anti-CD4 antibodies beginning three days before tumor implantation, followed by continued depletion at regular intervals to prevent CD4^⁺^ T cell recovery. (D) Kaplan-Meier survival analysis evaluating the impact of CD4^⁺^ T cell depletion on IL-12-armored CAR-T cell efficacy. C57BL/6J mice implanted with heterogeneous CT2A:CT2AvIII tumors were treated with EGFRvIII CAR scIL12 T cells in the presence or absence of anti-CD4 depletion antibodies. CD4^⁺^ T cell depletion did not significantly impact survival compared to isotype controls (ns, not significant), suggesting that CD4^⁺^ T cells are dispensable for IL-12-driven antitumor immunity in this model. (E) Experimental design of CD8^⁺^ T cell knock-out mouse studies. (F) Kaplan-Meier survival analysis demonstrating that IL-12-armored CAR-T cell efficacy is dependent on CD8^⁺^ T cells. Wild-type C57BL/6J mice and CD8^⁻^/^⁻^ mice were implanted with heterogeneous CT2A:CT2AvIII tumors and treated with EGFRvIII CAR scIL12 T cells. While EGFRvIII CAR scIL12 treatment significantly extended survival in C57BL/6J mice, no survival benefit was observed in CD8^⁻^/^⁻^ mice, confirming that CD8^⁺^ T cells are required for IL-12-mediated tumor clearance (p < 0.0001).

### CAR-T cells secreting IL-12 induce homeostatic mechanisms leading to persistence

To investigate how IL-12-armored CAR-T cells remodel the tumor immune landscape, we performed single-cell RNA sequencing (scRNA-seq) on CD45⁺ immune cells isolated from tumors treated with EGFRvIII CAR-T cells, EGFRvIII CAR-scIL12 T cells, or untreated controls 4 days post-treatment. To ensure our analyses focus on tissue-resident CD45 cells, we intravenously pre-treated mice with CD45 antibody prior to tissue harvesting and flow sorting to precisely remove any peripheral CD45^+^ immune cells. UMAP clustering (Fig. 3A) and annotation of cell clusters using reference atlases(20) identified immune and tumor cell populations (Fig. 3B, Supplemental Fig. 2). CAR-T cells were distinguished from endogenous T cells by MSGV retroviral long terminal repeat (LTR) expression, which demonstrated preferential CAR-T cell localization to clusters 1 and 5 (Fig. 3C). Flow cytometry quantification of CAR-T cells at multiple time points post CAR infusion confirmed a notable persistence of CAR⁺ T cells in EGFRvIII CAR-scIL-12 treated tumors compared to EGFRvIII CAR treated tumors (p < 0.01, Fig. 3D).

**Figure 3.**
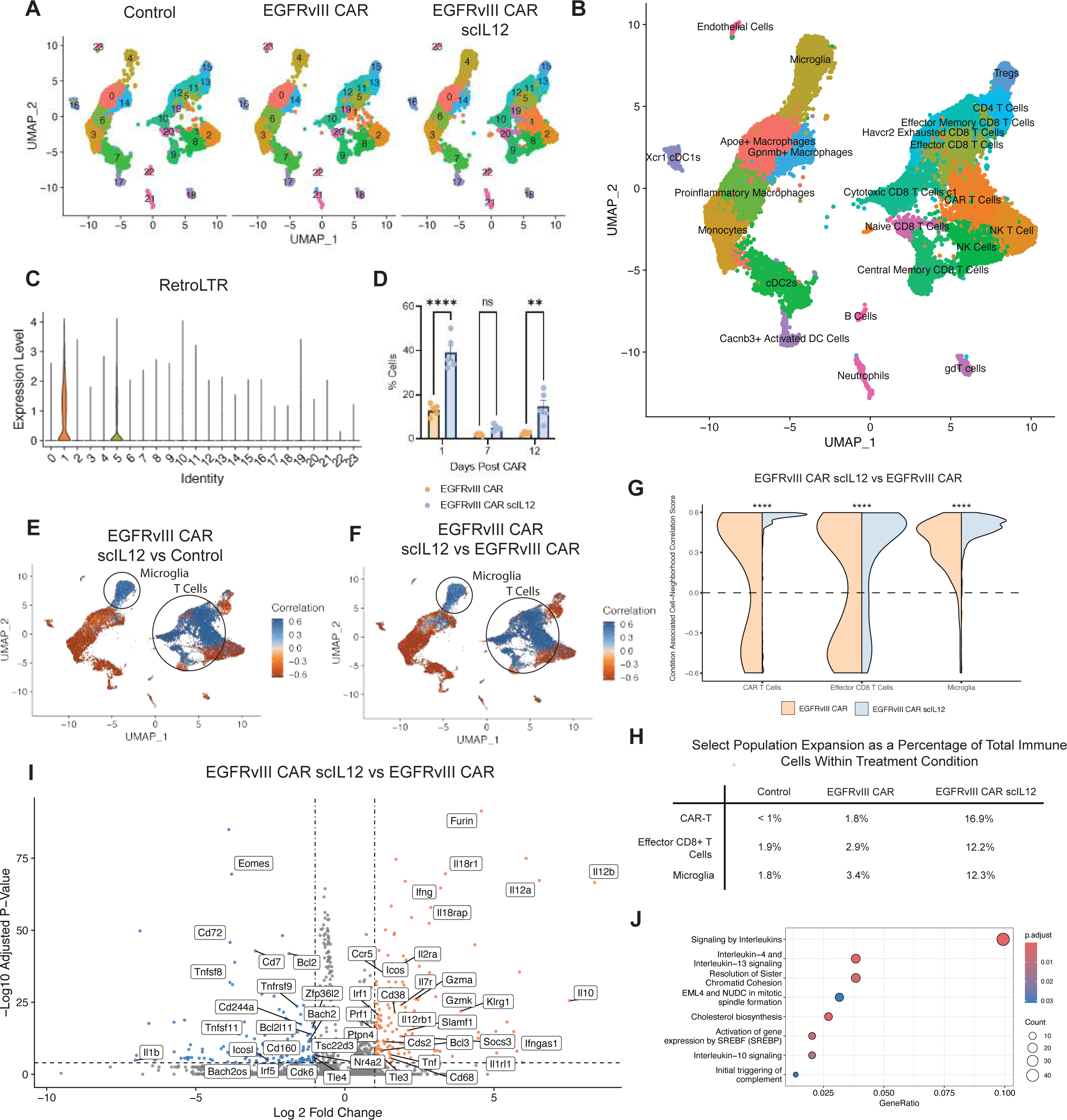
Single-Cell Transcriptomic Analysis of Tumor and Immune Cell Populations Following IL-12-Armored CAR-T Cell Therapy. (A) Uniform Manifold Approximation and Projection (UMAP) plot of immune and tumor cell populations from treated and control tumors. Single-cell RNA sequencing (scRNA-seq) was performed on CD45^⁺^ immune cells isolated from tumors of mice treated with EGFRvIII CAR-T cells, EGFRvIII CAR scIL12 T cells, or untreated controls. (B) Annotated UMAP plot identifying major cell types in the tumor microenvironment, including CD8+ T cells, macrophages, and dendritic cells. (C) Expression of the MSGV retroviral long terminal repeat (LTR) was used to track CAR-T cells. Analysis of the frequency of retroviral LTR integration site reads within UMAP clusters demonstrate distinct enrichment within few clusters. (D) Quantification of flow-cytometry validated CAR-T cells as a percentage of total T cells at various time intervals post-CAR-T cell treatment highlights significant expansions in CD8+ T cells in EGFRvIII CAR scIL12 treated groups (p<0.01). (E-F) Co-Varying Neighborhood Analysis (CNA) comparing cluster enrichment across treatment conditions. CNA analysis identified significant shifts in myeloid and T cell populations following IL-12-armored CAR-T cell treatment. (G) Condition-associated cell neighborhood correlation scores showing significant enrichment of CAR+ T cells, effector CD8^⁺^ T cells, and microglia in EGFRvIII CAR scIL12 T cell treated groups compared to EGFRvIII CAR-T cell treated groups. (H) Proportional expansion of key immune cell populations within treatment conditions, highlighting increased frequencies of effector CD8^⁺^ T cells, microglia, and CAR-T cells in EGFRvIII CAR scIL12-treated tumors compared to controls. (I) Differential gene expression analysis of EGFRvIII CAR scIL12 T cells versus standard EGFRvIII CAR-T cells. (J) Pathway enrichment analysis identifying major signaling pathways activated in IL-12-secreting CAR-T cells relative to CAR-T cells that do not secrete IL-12.

To further quantify treatment-induced immune shifts, we performed Co-Varying Neighborhood Analysis (CNA) (21) (Fig. 3E-F). CNA comparing EGFRvIII CAR scIL12 with EGFRvIII-CAR-T cells revealed significant changes in the representation of multiple immune cell types (Fig. 3F, Supplemental Fig. 3). The most pronounced changes in the IL-12 armored condition included increased CNA enrichment scores (Fig. 3G), and an overall rise in the proportion of CAR-T cells, endogenous effector CD8⁺ T cells, and microglia within the GBM immune infiltrate (Fig. 3H).

Differential gene expression analysis of EGFRvIII CAR scIL12 T-cells versus EGFRvIII CAR-T cells (Fig. 3I) provided deeper mechanistic insights into how IL-12 reshapes CAR-T cell function. EGFRv I I I CAR sc IL12 T-cells exhibited upregulation of activation-associated genes (Ifng, Il2ra, Prf1, Gzmb), survival genes (Bcl2), and T cell homing factors (Ccr5, Il7r). In contrast, genes associated with terminal exhaustion and differentiation (Eomes, Nr4a2) were downregulated.

Pathway enrichment analysis (Fig. 3J) identified several signaling cascades activated in EGFRv I I I CAR-sc IL-12 T cells, beyond canonical interleukin pathways. While the enrichment of IL-13 and IL-4 signaling pathways was at first surprising, recent studies have found these type 2 cytokines to alleviate CAR-T cell dysfunction and correlate with T-cell homeostasis and long-term persistence.(22) The enrichment of Sterol Regulatory Element-Binding Protein (SREBP) signaling and cholesterol biosynthesis pathways suggests that metabolic adaptation is crucial for CAR-T cell persistence and function in the brain.(23) Interestingly, these pathways have also been implicated as mediators of tumor cell metastases to the brain(24), potentially suggesting a mechanism of adaptation to the brain microenvironment. Cholesterol plays a critical role in T-cell synapse formation, lipid raft integrity, and sustained T-cell signaling, all of which are necessary for prolonged anti-tumor responses.(25)

### CAR-T cells Secreting IL-12 Reprogram the Tumor Microenvironment

To elucidate how IL-12 secretion by CAR-T cells reprograms the tumor microenvironment (TME), we analyzed changes in immune cell composition and intercellular signaling interactions. In addition to enriching the number of CAR-T cells, sc-RNAseq analysis revealed significant shifts across both the lymphoid and myeloid compartments following treatment with EGFRvIII CAR scIL12 T cells compared to treatment with EGFRvIII CAR-T cells (Fig. 3F, Supplementary Fig. 3).

Within the lymphoid compartment, IL-12 armoring significantly expanded endogenous CD8⁺ T cell subsets, including central memory, effector memory, and cytotoxic effector CD8⁺ T cells (Supplementary Fig. 3). Within the myeloid compartment, EGFRvIII CAR-scIL12 treatment selectively enriched microglia while reducing the proportion of macrophages and conventional dendritic cells (DCs) (Supplementary Fig. 3).

To further dissect how IL-12 contributes to immune remodeling, we performed CellChat (26) analysis to compare intercellular communication networks in EGFRvIII CAR-treated tumors vs. EGFRvIII CAR-scIL12-treated tumors. Analysis of differential interaction numbers (Fig. 4A) and strength (Fig. 4B) demonstrated that IL-12 armoring broadly enhanced intercellular communication within the TME. In EGFRvIII CAR-treated tumors (Fig. 4C), Apoe⁺ macrophages and proinflammatory macrophages were the primary sources of IL-12, with signaling effects largely restricted to central memory CD8⁺ T cells and Gpnmb⁺ macrophages. However, in EGFRvIII CAR scIL12-treated tumors (Fig. 4D), CAR-T cells unsurprisingly emerged as the dominant IL-12 producers, alongside newly identified IL-12-producing populations, including monocytes, microglia, and Gpnmb⁺ macrophages.

**Figure 4.**
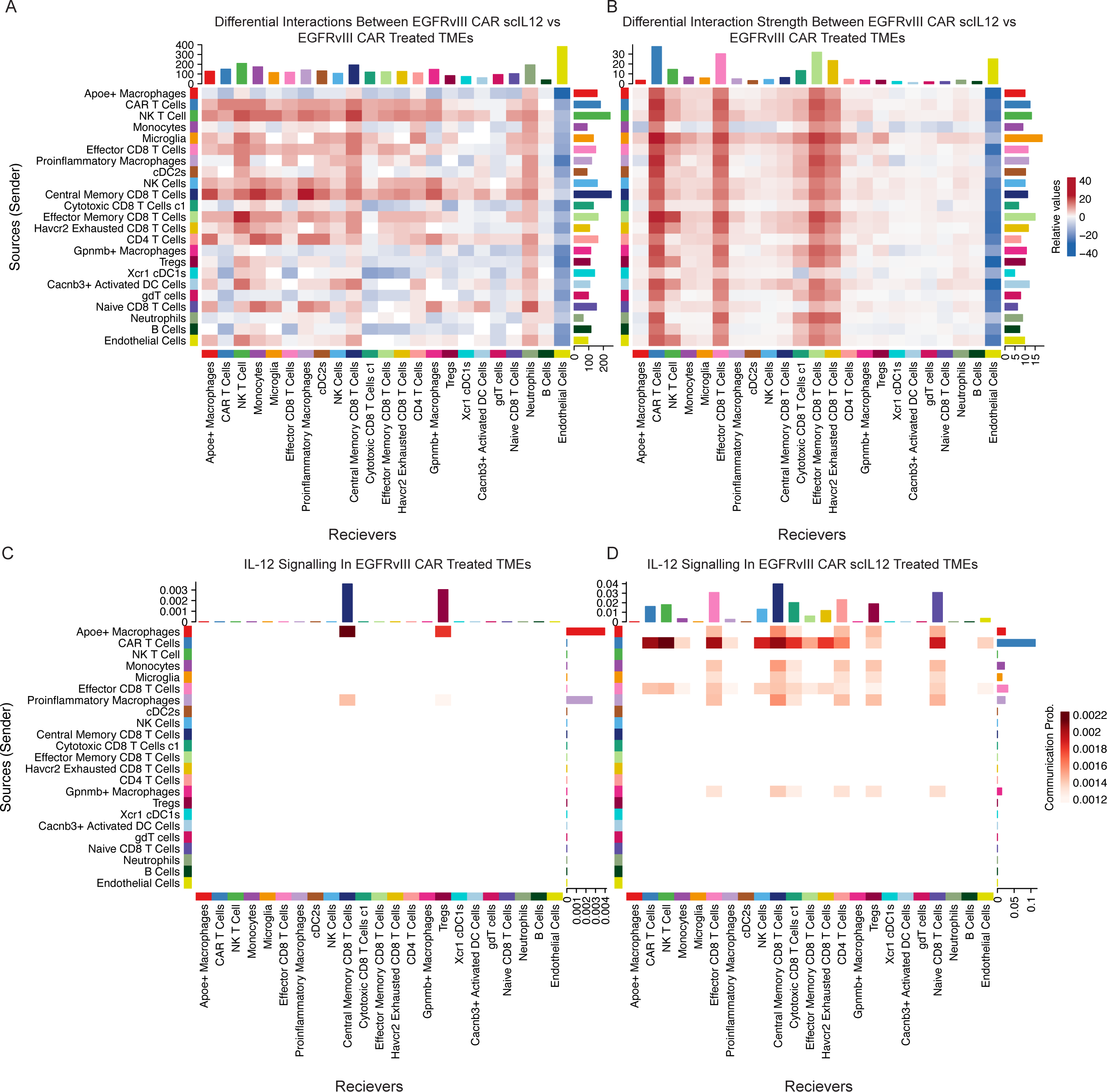
EGFRvIII CAR scIL12Remodel the Tumor Microenvironment to Enhance Antigen Presentation and T Cell Activation. (A) Single-cell RNA transcriptomic data were analyzed to determine shifts CellChat predicted intercellular interactions within the TME after treatment with EGFRvIII CAR or EGFRvIII CAR scIL12 T cells. The IL-12-armored CAR-T treatment significantly enhanced interactions among antigen-presenting cells (APCs), CAR-T cells, and endogenous effector CD8^⁺^ T cells. (B) The relative strength of intercellular communication was significantly increased between CAR-T cells, microglia, and effector CD8^⁺^ T cells in IL-12-treated tumors compared to controls. Proinflammatory macrophages, dendritic cells, and microglia emerged as key hubs of cellular communication in the IL-12-treated TME. (C) In standard EGFRvIII CAR-treated tumors, IL-12 signaling was largely restricted to Apoe^⁺^ macrophages and a limited subset of proinflammatory macrophages, with minimal engagement of adaptive immune cells. (D) In contrast, tumors treated with EGFRvIII CAR scIL12 T cells exhibited a significantly expanded IL-12 signaling network, with CAR-T cells, microglia, CD8^⁺^ T cells (effector, cytotoxic, and memory subsets), and NK cells all engaging in IL-12-driven interactions.

### IL-12-Armored CAR-T cells Recruit and Sustain a Polyclonal Endogenous T Cell Response

We hypothesized that IL-12 enhances survival with CAR-T cell therapy by recruiting and expanding endogenous T cells within the tumor microenvironment (TME), thereby broadening the repertoire of tumor-reactive T-cells. Our prior findings supported this hypothesis by demonstrating that EGFRvIII CAR scIL12-treated tumors exhibit increased T-cell infiltration (Fig. 3H). To further explore this effect, we performed TCR clonality analysis using VDJ region sequencing and stratifying CD4⁺ and CD8⁺ T-cell repertoires across treatment conditions.

To quantify T-cell repertoire diversity within the endogenous T-cell fraction, we excluded the CAR^+^ T-cells from analysis and calculated the Shannon entropy index, which measures overall TCR diversity, and the Gini index, which quantifies clonality skewness (Fig. 5A). Shannon entropy scores increased stepwise across treatment groups (Control < EGFRvIII CAR < EGFRvIII CAR scIL12), suggesting that IL-12 fosters the most diverse endogenous T cell response. However, the Gini index unexpectedly showed EGFRvIII CAR < Control < EGFRvIII CAR scIL12, indicating that standard CAR-T treatment had a more oligoclonal T cell population than untreated tumors, while IL-12 reversed this effect, fostering a more evenly distributed, polyclonal T cell repertoire. This seeming trend suggests that while standard CAR-T treatment induces expansion of tumor-reactive clones, it does so in a highly selective manner, restricting overall diversity. In contrast, IL-12 not only expands T cells but also prevents excessive clonal dominance, promoting a balanced and heterogeneous response, which may contribute to broader antigen recognition and epitope spreading.

**Figure 5.**
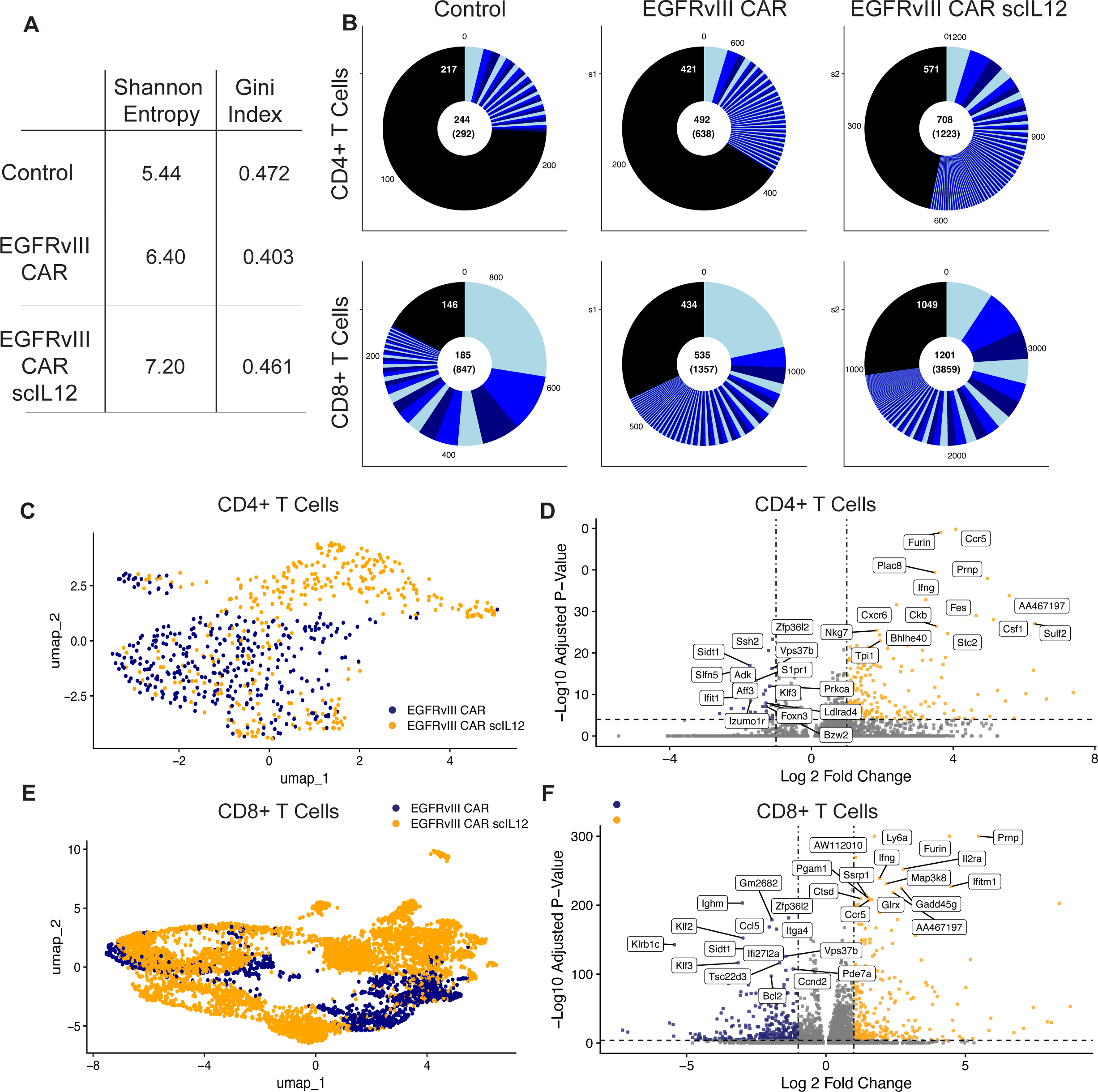
Clonal diversity and TCR repertoire analysis following EGFRvIII CAR-T cell therapy with and without IL-12 expression. (A) Shannon entropy index and Gini index analyses quantifying the diversity of T cell receptor (TCR) sequences across treatment groups within the non-adoptively transferred T cell compartment. (B) Donut charts displaying the relative proportions of expanded TCR clones within each treatment group stratified by endogenous CD4+ T cells and CD8+ T cells, illustrating how IL-12 expression promotes a broader TCR response compared to standard CAR-T cell therapy. Number in center of each donut plot represents number of unique TCR clonotypes in each condition whereas the parenthesized number in the center of each donut plot represents number of total numbers of TCR sequenced cells in each condition. (C-D) Endogenous CD4^⁺^ T cells in the IL-12 group clustered separately from those in the standard CAR-T group, with transcriptional profiles indicative of enhanced activation and differentiation into helper and effector subsets. (E-F) Endogenous CD8^⁺^ T cells in the IL-12 treatment group clustered away from standard CAR-T-treated CD8^⁺^ T cells, suggesting functional reprogramming.

Donut plots of TCR expansion (Fig. 5B) further illustrated these findings, with EGFRvIII CAR-treated tumors displaying a less polyclonally expanded TCR landscape, whereas EGFRvIII CAR scIL12-treated tumors exhibited greater polyclonal expansion within both endogenous CD4⁺ and CD8⁺ compartments.

To determine how IL-12 influences endogenous T cell function, we performed UMAP clustering of endogenous CD4⁺ and CD8⁺ T cells from heterogeneous tumors treated with either EGFRvIII CAR or EGFRvIII CAR scIL12 (Fig. 5D, 5F). While a subset of CD4⁺ T cells from EGFRvIII CAR-scIL12-treated tumors clustered similarly to those from EGFRvIII CAR-treated mice, an additional, transcriptionally distinct CD4⁺ T cell population emerged exclusively in the IL-12 condition, suggesting functional reprogramming (Fig. 5C). Differential gene expression analysis (Fig. 5D) identified significant upregulation of IFNγ, Gzmb, and Ccr5, indicative of an enhanced effector phenotype with increased tumor-homing potential in CD4⁺ T cells from EGFRvIII CAR-scIL-12 treated tumors. A similar effect was observed in CD8⁺ T cells, where UMAP clustering (Fig. 5E) revealed a clear transcriptional shift in IL-12-treated tumors. Differential gene expression analysis (Fig. 5F) identified upregulation of Il2ra, Ifng, and Prf1, suggesting enhanced cytokine responsiveness and cytotoxic potential.

Interestingly, Ccr5, Furin, and Ifng were all upregulated in both endogenous CD8⁺ and CD4⁺ T cells in EGFRvII CAR-IL12 treated tumors, suggesting a shared transcriptional program that enhances immune coordination. Ccr5 upregulation indicates enhanced tumor homing and infiltration, as Ccr5 is a key chemokine receptor involved in T cell trafficking to inflamed tissues.(27–29) Recent findings indicate that a Th1-specific super-enhancer region upstream of the Furin gene enhances IFNγ secretion, reinforcing a proinflammatory microenvironment that supports durable immune activation against tumors (30, 31). Ifng upregulation in both CD4⁺ and CD8⁺ T cells reinforces a highly proinflammatory TME, likely further amplifying antigen presentation and effector function.

### CAR-T cells Secreting IL-12 Reprogram Microglia to Create a Proinflammatory Microenvironment

Traditionally, myeloid cells in the tumor microenvironment have been considered a major barrier to immunotherapy efficacy (32–34). However, despite the significant increase in brain-resident microglia in the EGFRvIII CAR-scIL12 condition, we observed enhanced survival, suggesting that IL-12 may be altering the phenotype of microglia to support endogenous adaptive immune responses.

Within the microglial compartment, EGFRvIII CAR scIL12 cell therapy led to significant upregulation of antigen presentation genes (H2-DMa, H2-Ab1, H2-Aa) (Fig. 6A), indicating enhanced antigen processing capacity. Additionally, microglia from IL-12-treated tumors exhibited increased expression of interferon response genes (Stat1, Irf1). Analysis of conventional dendritic cells (cDC1s) revealed increased expression of survival and metabolic genes (Serpinb9b, Pla2g15) in EGFRv III CAR sc IL12-treated tumors (Fig. 6B). Similarly, macrophages in the EGFRvIII CAR scIL12 condition exhibited upregulation of Ly6a (Sca-1) and Ifi204, genes linked to cell activation and interferon responses (Fig. 6C).

**Figure 6.**
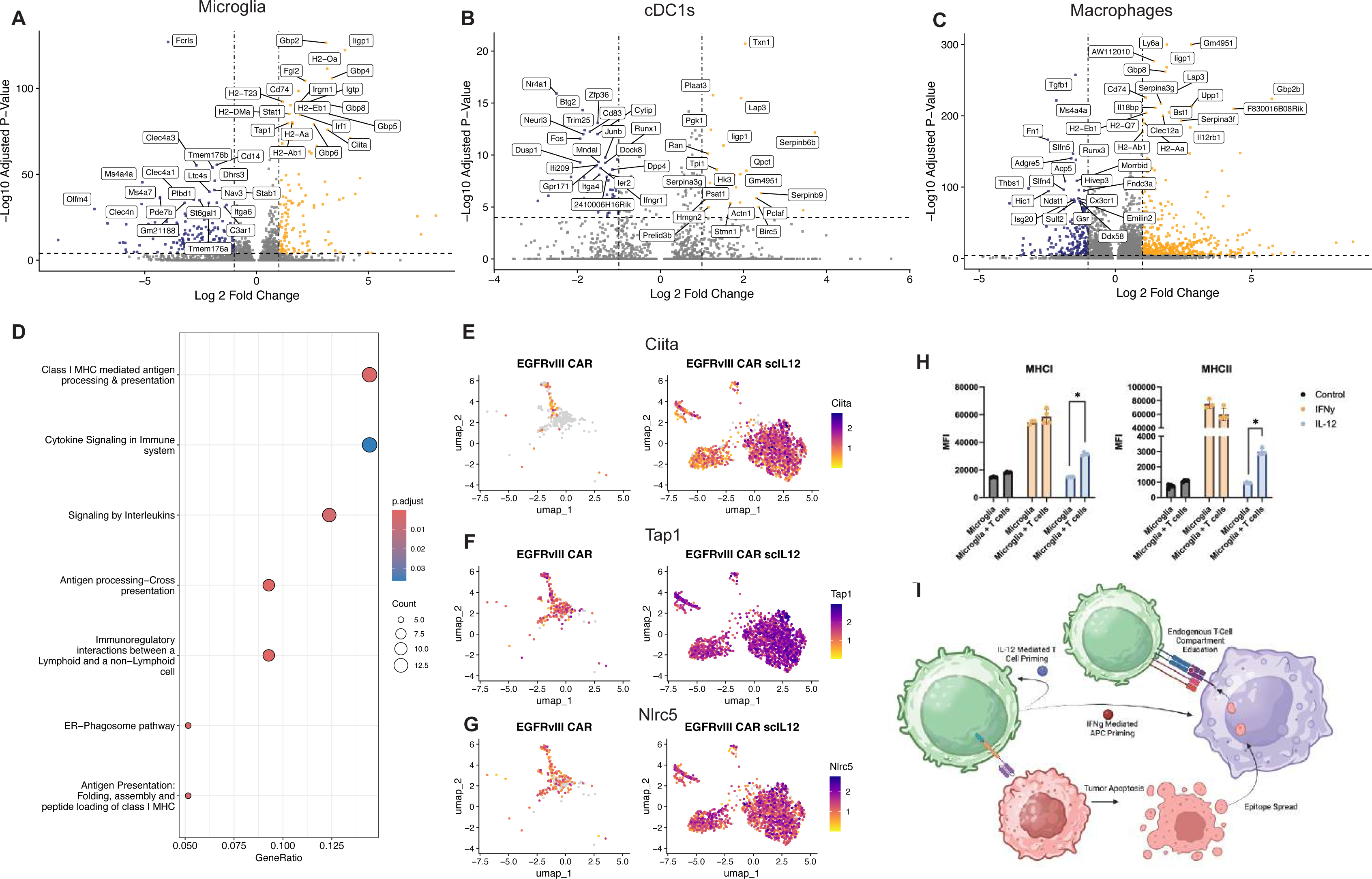
EGFRvIII CAR scIL12Reprogram Microglia to Enhance Antigen Presentation and Immune Activation. (A) Differential gene expression in microglia following EGFRvIII CAR-T cell therapy with or without IL-12. Volcano plot highlighting genes upregulated in EGFRvIII CAR scIL12-treated tumors, revealing significant enrichment in genes involved in antigen presentation (H2-Ab1, H2-DMa, Ciita, Tap1, Nlrc5), interferon signaling (Stat1, Irf1), and immune activation (Gbp2, Cd74) (B) Differential gene expression in conventional dendritic cells (cDC1s). cDC1s from tumors treated with IL-12-secreting CAR-T cells displayed upregulation of survival and metabolic genes (Serpinb9b, Pla2g15), suggesting enhanced functionality and longevity of antigen-presenting cells in the IL-12 treatment group. (C) Differential gene expression in macrophages following CAR-T treatment. (D) Pathway enrichment analysis of microglia from tumors treated with EGFRvIII CAR scIL12 T cells. Gene set enrichment analysis identified class I and II MHC antigen presentation, cytokine signaling, and cross-presentation pathways as significantly enriched in microglia from IL-12-treated tumors, suggesting that microglia actively contribute to antigen processing and T cell priming. (E-G) UMAP projection of key antigen presentation genes (Ciita, Tap1, Nlrc5) in microglia. Expression of Ciita, Tap1, and Nlrc5 was significantly upregulated in microglia from IL-12-treated tumors, confirming that IL-12 promotes antigen-presenting capacity in microglia. (H) Bar graphs depicting MHC-II and MHC-I expression levels across treatment groups in IMG mouse microglial cell lines cultured in vitro with interferon-gamma (IFNg) or IL-12, demonstrating significant upregulation of MHC molecules when cells are in the presence of IFNg or both T cells and IL-12. (I) Schematic summarizing the proposed model of IL-12-mediated microglial reprogramming. IL-12 secretion by CAR-T cells promotes a proinflammatory phenotype in microglia mediated via INFg, enhancing their ability to process and present antigens to endogenous T cells, thus facilitating epitope spreading and sustained immune activation.

Pathway enrichment analysis identified MHC class I-mediated antigen processing and presentation as significantly enriched in microglia from EGFRvIII CAR-IL-12-treated tumors (Fig. 6D). Given the observed expansion of microglial populations, we hypothesized that IL-12-induced microglial activation facilitates T cell recruitment and maintenance. Supporting this, key antigen presentation regulators (Ciita, Tap1, and Nlrc5) were significantly upregulated in the microglia of EGFRvIII CAR-scIL12 treated mice (Fig. 6E-G), reinforcing the role of IL-12 in promoting antigen presentation and T-cell priming.

To determine whether IL-12 alone is sufficient to drive MHC-I and MHC-II expression in microglia, we cultured IMG microglial cell lines in the presence of IL-12, naïve murine T-cells, or both and analyzed MHC c lass I and I I s urface expression by f low c ytometry (Fig. 6H). IFNγ served as a positive control, given its well-characterized role in MHC upregulation. (35–38) While IL-12 alone failed to induce significant MHC-I or MHC-II expression, co-culturing IL-12-treated microglia with naïve T cells significantly enhanced MHC expression (Fig. 6H).

Based on these findings, we propose a model in which IL-12 secreted by EGFRv III CAR sc IL 12 T-cells enhances T-cell activation and IFNγ secretion, which in turn reprograms microglia to adopt a proinflammatory, antigen-presenting phenotype (Fig. 6I). This feedback loop amplifies T cell recruitment, persistence, and possible epitope spreading, likely explaining the superior tumor clearance and prolonged survival observed in IL-12-treated glioblastoma models.

## DISCUSSION

The failure of current treatments in GBM, including CAR-T cell immunotherapy, stems from two interrelated challenges: (i) tumor heterogeneity, which allows antigen-negative clones to evade targeted therapies, and (ii) an immunosuppressive microenvironment, which limits endogenous immune responses. In this study, we demonstrate that EGFRvIII-targeted CAR-T cells armored with single-chain IL-12 (scIL12) address both of these challenges through a dual mechanism of action that involves direct tumor targeting and microenvironmental reprogramming. Our results show that IL-12 secretion obviates the need for lymphodepletion, enhances CAR-T cell persistence, and engages endogenous CD8⁺ T cells to clear antigen-negative tumor cells. These findings position IL-12-armored CAR-T cells as a powerful strategy to extend the efficacy of CAR-T therapy to GBM and potentially other solid tumors.

We found that conventional EGFRvIII-targeted CAR-T cells fail to control mixed EGFRvIII+/EGFRvIII− tumors, reflecting clinical observations where antigen escape limits therapeutic durability. Our data reveal that IL-12 armoring circumvents this challenge by engaging endogenous CD8+ T cells to eliminate antigen-negative tumor cells through epitope spreading. This mechanism was strictly dependent on CD8+ T cells, as depletion experiments in CD8−/− mice abolished therapeutic efficacy, whereas NK cell depletion had no impact. These findings align with studies highlighting IL-12’s role in enhancing traditional antigen-presenting cell cross-presentation (15) but extend this paradigm by uniquely implicating microglia—the brain’s resident myeloid compartment—as critical antigen-presenting cells (APCs) in the GBM microenvironment.

A critical observation from our study was that IL-12 alone was insufficient to fully induce this phenotype in microglia; instead, the presence of T cells was required for optimal antigen presentation capacity. In vitro experiments demonstrated that IL-12 treatment alone did not significantly upregulate MHC-I or MHC-II expression in microglial cell lines, but co-culture with naïve T cells resulted in increased expression of antigen presentation machinery. This suggests a cooperative model in which IL-12 initiates myeloid activation, while T cell-derived cytokines such as IFNγ reinforce the antigen-presenting function of microglia. Given the relative paucity of T cells in human GBM samples, our findings further support the importance of exogenously administered CAR-T cells as an initiator of the feed forward loops that result in T cell accumulation and epitope spreading.

Beyond engaging endogenous immunity, IL-12 secretion fundamentally enhances CAR-T cell fitness within the immunosuppressive GBM microenvironment. scRNA-seq revealed that IL-12-armored CAR-T cells exhibited prolonged persistence in tumors, with significant enrichment of cytotoxic (*Ifng*, *Gzmb*, *Prf1*) and homing (*Ccr5*) genes compared to conventional CAR-T cells. Longitudinal tracking demonstrated that CAR-T cells secreting IL-12 remained detectable in tumors for up to 12 days post-infusion, whereas unarmored CAR-T cells were largely eliminated by day 4. This enhanced persistence correlated with upregulation of pro-survival pathways and suppression of exhaustion-associated markers. Our findings confirm IL-12’s role in enhancing T cell activity and resistance to exhaustion in solid tumors (39–41).

A striking finding of this study is that IL-12 armoring eliminates the need for lymphodepletion, a standard preconditioning step associated with significant toxicity. In our study, while conventional CAR-T cells required total body irradiation (TBI) to control homogeneous tumors, IL-12-armored CAR-T cells achieved 100% survival *without preconditioning*. Conversely, lymphodepletion abrogated efficacy in heterogeneous models by depleting endogenous lymphocytes necessary for epitope spreading. Clinically, circumventing lymphodepletion could expand eligibility for GBM patients with poor performance status that would preclude lymphodepletion, reduce treatment-related complications such as cytopenias and infections, and help lower the economic burden associated with lymphodepletion regimens in CAR-T cell therapy.

The success of IL-12-armored CAR-T cells in preclinical models presents key challenges for clinical application. While IL-12’s pleiotropic effects enhance efficacy, its systemic toxicity—such as cytokine release syndrome (CRS) and neuroinflammation—must be carefully managed. Landmark studies have demonstrated that systemic IL-12 administration can lead to severe toxicity, limiting its therapeutic potential.(42) Strategies including inducible expression systems (e.g., tetracycline-regulated promoters or NFAT regulated systems)(43, 44) or localized intracranial delivery have been proposed to mitigate these risks. Recent data have suggested that intracranial delivery of IL-12, may in fact be neuroprotective (45) and has been safely and repeatedly delivered locally into the brain, including in several ongoing clinical trials for the treatment of GBM [NCT05095441, NCT02062827].

Future work should focus on elucidating how IL-12 mechanistically enhances antigen presentation and endogenous T-cell recruitment within the TME. Specifically, understanding its role in microglial polarization and antigen presentation may reveal new therapeutic targets for GBM CAR-T cell optimization. Additionally, refining IL-12 delivery strategies to balance efficacy and toxicity will be essential for translating these insights into clinical applications. Surface-tethering approaches, in which IL-12 is localized to the CAR-T cell membrane, may represent a feasible next step to mitigate toxicity while preserving local efficacy (46, 47). Furthermore, directed evolution approaches to engineer IL-12 variants with reduced toxicity while maintaining potency could enhance clinical viability (48).

In summary, we demonstrate that IL-12-armored CAR-T cells represent a multifaceted therapy capable of addressing the dual challenges of antigen heterogeneity and immunosuppression in GBM. By enhancing CAR-T cell persistence, upregulating antigen presentation machinery on microglia, and recruiting polyclonal endogenous T-cells, this strategy achieves durable tumor control without lymphodepletion. With further refinement, IL-12-secreting CAR-T cells could transform the treatment landscape for GBM and other immunologically “cold” malignancies.

## MATERIALS AND METHODS

### Mouse Models

6–10 week old female mice were used unless otherwise specified. C57BL/6J mice were obtained from Charles River Laboratories (Wilmington, MA). Transgenic strains including B6.129S2-Cd8atm1Mak/J (CD8⁻/⁻; JAX Strain #002665), B6.129S2-Tcratm1Mom/J (TCRα⁻/⁻; JAX Strain #002116), and B6.129S7-Rag1tm1Mom/J (RAG1⁻/⁻; JAX Strain #002216) mice were acquired from Jackson Laboratory (Bar Harbor, ME). All animal experiments were conducted following protocols approved by the Duke University Institutional Animal Care and Use Committee (IACUC).

### Glioma Cell Line Culture and Implantation

CT2A murine glioma cells (Millipore Sigma SCC194) and CT2AvIII cells (lentiviral EGFRvIII-engineered) were maintained in DMEM (Gibco) supplemented with 10% FBS, 2 mM L-glutamine, and 4.5 mg/mL glucose. For orthotopic implantation, cells were resuspended in PBS (4×10D cells/mL) mixed 1:1 with 3% methylcellulose (Sigma) to prevent aggregation and stereotactically injected into the right basal ganglia to deliver a total of 5×10D cells (2 mm lateral to bregma, 4 mm depth). CAR-T cells (2×10D) were administered intracranially 7 days post-tumor engraftment. Mice were randomized into treatment groups at the time of tumor implantation. Potential confounders such as cage position or experiment order were not formally controlled. Mice were monitored daily for survival.

### Generation and Engineering of CAR Constructs

The third-generation EGFRvIII-specific CAR was constructed using the MSGV1 retroviral backbone (49). For EGFRvIII CAR scIL12 T cells, the construct was modified to include an encephalomyocarditis virus (EMCV) internal ribosome entry site (IRES) followed by a single-chain murine IL-12 gene, generated by linking the p35 (Il12a) and p40 (Il12b) subunits with a (G₄S)₃ flexible peptide linker. Retroviral supernatant was produced by co-transfecting HEK 293T cells (ATCC CRL-3216) with the CAR plasmid and pCL-Eco packaging plasmid (Novus Biologicals, #NBP2-29540) using Lipofectamine 2000 (Invitrogen, #11668019). Viral supernatant was harvested 24 hours post-transfection, filtered (0.45 μm), and used immediately. Murine splenocytes were isolated from C57BL/6 mice and activated with 2 μg/mL Concanavalin A (InvivoGen, #inh-cona-2) and 50 IU/mL recombinant murine IL-2 (PeproTech, #212-12) for 48 hours.

Activated T cells were transduced on Retronectin (Takara Bio, #T100A)-coated plates via spinoculation (2,000 × g, 90 minutes, 32°C) with viral supernatant. Transduced cells were expanded for 7 days in RPMI-1640 supplemented with 10% FBS and 50 IU/mL IL-2.

### Tissue Harvesting and Tumor Dissociation

Mice were transcardially perfused with PBS + 10 IU/mL heparin. Intracranial tumors were harvested and weighed. Tumor-bearing hemispheres were dissociated in digestion buffer (HBSS + 2.5 mg/mL Liberase TL [Sigma, #5401020001], 2.5 mg/mL Liberase DL [Sigma,#54602020001], and 100 U/mL DNase I [Roche, #10104159001]) using a Dounce homogenizer. Samples were incubated at 37°C for 30 minutes, quenched with HBSS, and filtered through 70 μm cell strainers. Tumor-infiltrating lymphocytes (TILs) were isolated using a 30% isotonic Percoll (Sigma, #P1644) gradient (500 × g, 30 minutes, 18°C), followed by red blood cell lysis with ACK buffer (Gibco, #A1049201).

### Flow Cytometry and Immune Cell Profiling

Single-cell suspensions from tumors were stained with LIVE/DEAD Aqua Viability Dye (Thermo Fisher, #L34957) for 20 minutes at 4°C. Surface staining was performed using fluorophore-conjugated antibodies. For intracellular staining, cells were fixed and permeabilized using the Foxp3/Transcription Factor Staining Buffer Set (eBioscience, #00-5523-00). Absolute cell counts were determined using CountBright Absolute Counting Beads (Thermo Fisher, #C36950). Compensation was performed using UltraComp eBeads (Thermo Fisher, #01-2222-42) and ArC Amine Reactive Compensation Beads (Thermo Fisher, #A10346). Data were acquired on a BD LSRFortessa (BD Biosciences) and analyzed using FlowJo v10.8 (Tree Star).

### In Vitro Cytotoxicity Assessment of CAR-T cells

CAR-T cell cytotoxicity was assessed by co-culturing effector cells with target tumor cells labeled with 5 μM CFSE (Thermo Fisher, #C34554). Cells were plated at effector-to-target (E:T) ratios of 0.25:1 to 4:1 in 96-well U-bottom plates. After 24 hours at 37°C, cells were washed and analyzed by high-throughput flow cytometry (BD Fortessa). Tumor cell lysis was quantified as the percentage of viable cells remaining within the CFSE+ population.

### Quantification of Cytokine Secretion

Supernatant IL-12 levels were quantified using the Mouse IL-12 (p70) ELISA Kit (Invitrogen, #BMS616). CAR-T cells (2×10D/mL) were cultured in serum-free RPMI-1640 for 24 hours, and supernatants were assayed in triplicate per the manufacturer’s protocol.

### Histopathological Analysis and Immunohistochemistry

Brains were fixed in 4% paraformaldehyde (PFA; Sigma, #P6148) for 48 hours, paraffin-embedded, and sectioned at 4 μm thickness. For IHC, sections were deparaffinized, subjected to antigen retrieval (10 mM citrate buffer, pH 6.0), and incubated overnight at 4°C with anti-EGFRvIII monoclonal antibody (clone L8A4; Sigma, #MABC1126; 1:200). Detection was performed using Goat Anti-Mouse IgG-HRP (Sigma, #AP308P; 1:500) and DAB substrate (Vector Laboratories, #SK-4100). Sections were counterstained with hematoxylin (Sigma, #H9627).

### In Vivo Depletion of CD4+ T Cells and NK Cells

For CD4+ T cell depletion, mice received intraperitoneal (i.p.) injections of 200 μg anti-CD4 (clone GK1.5; BioXCell, #BE0003-1) or isotype control (clone LTF-2; BioXCell, #BE0090) in PBS three days prior to tumor implantation, followed by 200 μg doses on day 2 post-implantation and every three days thereafter. For NK cell depletion, mice were injected i.p. with 500 μg anti-NK1.1 (clone PK136; BioXCell, #BE0036) or isotype control (clone C1.18.4; BioXCell, #BE0085) two days before tumor implantation, followed by 250 μg doses one day prior to implantation and every three days.

### Microglia Stimulation and Antigen Presentation Assays

Immortalized mouse microglia (IMG cells; RRID: CVCL_B5IN) were cultured alone or with isolated 1×10D murine splenic T cells (Miltenyi Biotec, #130-095-130) in RPMI-1640 + 10% FBS. Cultures were untreated or stimulated with 10 μg/mL recombinant murine IFNγ (PeproTech, #315-05) or IL-12 (PeproTech, #210-12) for 24 hours. Cells were stained with anti-MHC class and MHC class II antibodies and analyzed by flow cytometry as described above.

### Single-Cell Transcriptomic and TCR Sequencing

For *in vivo* labeling, mice received intravenous (i.v.) injections of 2 μg PE-conjugated anti-CD45 (clone 30-F11; BioLegend, #103105) 3 minutes prior to euthanasia. Brains were harvested, dissociated, and stained with LIVE/DEAD Aqua and APC-conjugated anti-CD45 (clone 30-F11; BioLegend, #103111). CD45-APC+, CD45-PE-, live cells were sorted on a FACSAria II (BD Biosciences) into chilled FBS and processed immediately. Single-cell libraries were prepared using the Chromium Next GEM Single Cell 5′ Kit (10X Genomics, #1000263) and sequenced on an Illumina NovaSeq S4 (PE150).

### Bioinformatic Analysis of Single-Cell Data

Raw sequencing data were processed using Cell Ranger v3.0.2 (10X Genomics) to generate gene expression matrices. Data were imported into Seurat v4.2 for quality control (QC), normalization, and clustering. Low-quality cells (<200 genes/cell, >10% mitochondrial reads) were excluded. The top 3,000 variable genes were used for principal component analysis (PCA), and the first 50 PCs were used for UMAP visualization. Cell types were annotated using scType (20) and manual curation based on canonical markers. Co-varying neighborhood analysis (CNA) was performed to compare cluster abundance across conditions (21). Cell-cell communication networks were inferred using CellChat v1.0.0 (26).

### TCR Clonality and Repertoire Analysis

TCR clonotypes were identified using Cell Ranger V(D)J (10X Genomics) and mm10 reference genome. Only productive, in-frame TCRα and TCRβ chains were considered. Dominant CDR3β sequences defined clonotypes, and clonal expansion was analyzed using Platypus v3.1 (50). TCR clonotype diversity was assessed using Shannon Entropy and the Gini Index. Both metrics were calculated using the vegan and DescTools R packages.

### Statistical Methods and Data Analysis

Survival curves were analyzed using Wilcoxon signed-rank test. Normality was assumed; non-parametric tests (Mann-Whitney, Kruskal-Wallis) were used when appropriate. Flow cytometry and cytokine data were compared using unpaired two-tailed t test, Mann Whitney U, or Kruskal-Wallis test. *p < 0.05* was considered significant. Analyses used GraphPad Prism v9.3.1 and R v4.2.2.

## ACKNOWLEDGEMENTS

We express special thanks to Dr. Simon Gregory, Karen Abramson, Stephanie Arvai, and all the other staff at the Molecular Genomics Core at Duke University. Furthermore, we thank David Snyder and Duke University’s Division of Laboratory Animal Resources (DLAR) team for their dedication to humane care and use of research animals used in our work comprised in this manuscript.

## DECLARATIONS

### Competing Interests

The authors have declared that no conflict of interest exists.

### Data availability Statement

Data are available upon reasonable request.

### Contributions

S.H.S. and A.A.M: Experimental design, experiments, data acquisition, analysis, drafting, revision. K.M.H., S.L.C., K.K.P., E.M., A.M.S., D.W.: Experiments, data analysis. C.M.S., P.E.F., L.S.-P.: Technical expertise, experimental design, manuscript review. J.H.S., A.P.P.: Experimental design, supervision, funding, manuscript editing/approval.

## Notes

### Competing Interest Statement

The authors have declared no competing interest.

