## Supplementary Figures for "IL-12 Secreting CAR-T Cells Reprogram the Tumor Microenvironment and Improve Efficacy Against Heterogeneous Models of Glioblastoma"

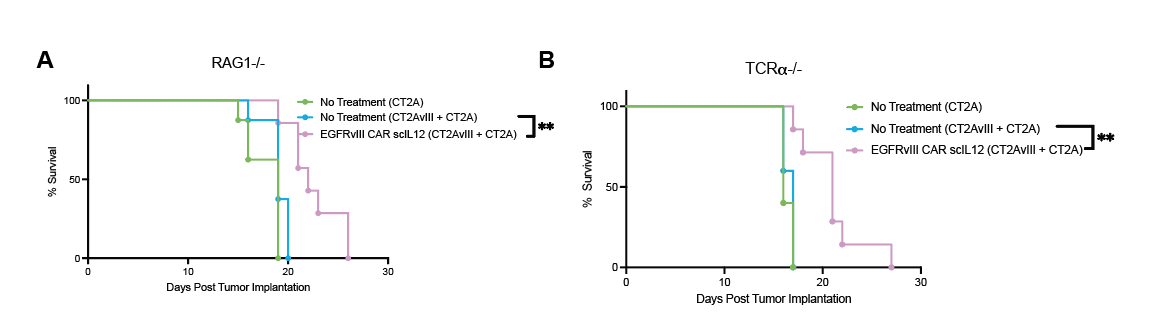

**Supplemental Figure 1: IL-12-Armored CAR-T Cells Require an Intact Adaptive Immune System for Efficacy in Heterogeneous Tumors**

(A) Kaplan-Meier survival curves for RAG1⁻/⁻ mice implanted with CT2A or CT2AvIII + CT2A tumors and treated with EGFRvIII CAR-scIL-12 T cells. RAG1⁻/⁻ mice, which lack mature T and B cells, failed to achieve long-term survival following IL-12-armored CAR-T cell treatment. (B) Kaplan-Meier survival curves for TCRα⁻/⁻ mice implanted with CT2A or CT2AvIII + CT2A tumors and treated with EGFRvIII CAR-scIL-12 T cells. TCRα⁻/⁻ mice, which lack functional αβ T cells (including CD4⁺ and CD8⁺ T cells), also exhibited a failure to control tumor progression, reinforcing the critical role of T cell-mediated immunity in IL-12-driven tumor clearance. Statistical significance is indicated by asterisks (*p < 0.05, **p < 0.01).

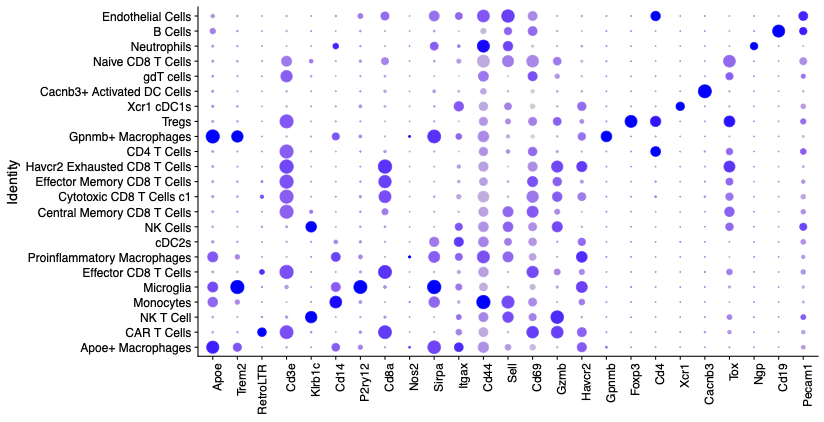

**Supplemental Figure 2: Single-Cell Gene Expression Analysis of Tumor-Infiltrating Immune Populations Following CAR-T Cell Therapy.**

Annotation dot plot displays the expression of key immune and lineage-defining markers across various tumor-infiltrating immune populations following treatment with EGFRvIII CAR-T cells and IL-12-armored CAR-T cells. Rows represent distinct immune cell subsets, and columns represent selected marker genes associated with T cells, myeloid cells, and antigen-presenting cells (APCs). CAR-T cells express LTR (retroviral integration marker) and Cd3e. CD8⁺ T cells (Effector, Central Memory, and Exhausted subsets) express Cd8a, Gzmb, and Cd69, with increased activation markers in effector populations. CD4⁺ T cells express Cd4 and regulatory T cell markers (Foxp3, Havcr2) within the TME. Microglia and macrophage subsets display distinct expression profiles, with P2ry12 and Trem2 marking microglia, while Cd14, Nos2, and Sirpa indicate activated macrophages. Dendritic cells (cDC1s, cDC2s) express Xcr1, Itgax, and Cacnb3. NK and NK-T cells show expression of Nkg7 and Gzmb, indicating cytotoxic function. Apoe⁺ macrophages, previously linked to an immunosuppressive phenotype, show distinct expression signatures compared to proinflammatory macrophages.

**Supplemental Figure 3: Condition-Associated Cell-Neighbor Correlation Scores Following EGFRvIII CAR and EGFRvIII CAR scIL12 Therapy**

(A) Violin plots depicting the condition-associated cell-neighborhood correlation scores comparing EGFRvIII CAR scIL12-treated samples to control-treated immune infiltrate in the tumor microenvironment. (B) Violin plots of the condition-associated cell-neighborhood correlation scores comparing EGFRvIII CAR scIL12-treated samples to EGFRvIII CAR-treated immune infiltrate in the tumor microenvironment. Significant enrichment of CAR⁺ T cells, effector CD8⁺ T cells, and microglia is observed in EGFRvIII CAR scIL12-treated groups, suggesting a distinct immune microenvironment polarization associated with IL-12 expression. Other immune subsets, including regulatory T cells (Tregs), dendritic cell populations (cDC1s, cDC2s), and macrophages, exhibit differential neighborhood correlations across treatment conditions. Statistical significance is indicated by asterisks (*p < 0.05, **p < 0.01, ***p < 0.001, ****p < 0.0001).
